# Genetic manipulation of stress-induced mitogen-activated protein kinase modulates early stages of the nodulation process in *Medicago sativa*

**DOI:** 10.1101/2022.11.09.515844

**Authors:** Kateřina Hlaváčková, Olga Šamajová, Miroslava Hrbáčková, Jozef Šamaj, Miroslav Ovečka

## Abstract

Leguminous plants have established a mutualistic endosymbiotic interaction with nitrogen-fixing rhizobia to secure nitrogen sources in new specialized organs called root nodules. Before nodule formation, the development of early symbiotic structures is essential for rhizobia docking, internalization, targeted delivery, and intracellular accommodation. We have recently reported that overexpression of stress-induced mitogen-activated protein kinase (SIMK) in alfalfa affects root hair, nodule and shoot formation. However, detailed subcellular spatial distribution, activation, and developmental relocation of SIMK during the early stages of alfalfa nodulation remain unclear. Here, we qualitatively and quantitatively characterized SIMK distribution patterns in rhizobium-infected root hairs using live-cell imaging and immunolocalization, employing alfalfa stable transgenic lines with genetically manipulated SIMK abundance and kinase activity. In the *SIMKK-RNAi* line, showing downregulation of *SIMKK* and *SIMK*, we found a considerably decreased accumulation of phosphorylated SIMK around infection pockets and infection threads, which was strongly increased in the GFP-SIMK line, constitutively overexpressing GFP-tagged SIMK. Thus, genetically manipulated SIMK modulates root hair capacity to form infection pockets and infection threads. These results shed new light on SIMK spatio-temporal participation in the early interactions between alfalfa and rhizobia, and its internalization into root hairs, showing that local accumulation of active SIMK indeed modulates nodulation in alfalfa.

**One sentence summary:** Genetic manipulation of SIMK in alfalfa revealed that SIMK modulates root hair capacity to form infection pockets and infection threads during the early interactions between alfalfa and rhizobia.

## Introduction

Nitrogen shortage in the soil is one of the major factors restricting the growth and productivity of plants, including crops. To overcome or alleviate this limitation, *Medicago sativa* L. (alfalfa), a legume crop of high agronomic and ecological importance, is able to acquire nitrogen by symbiotic hosting of nitrogen-fixing rhizobia in *de novo* formed specialized organs, root nodules (Checcucci *et al*., 2016; Wang *et al*., 2018). Root nodules provide rhizobia with favorable conditions to convert atmospheric dinitrogen (N_2_) into ammonia (NH_3_) in the process of biological nitrogen fixation. Rhizobia export N-rich compounds to the host plant in exchange for carbohydrates that are utilized by rhizobia as a source of carbon and energy (White *et al*., 2007; Oldroyd *et al*., 2011). The legume-rhizobium symbiosis is established through a complex developmental process that starts with the exchange of signaling molecules between the host and symbiont, and the activation of signal transduction pathways, triggering the nodulation program in the host legume plant (Oldroyd, 2013; Yang *et al*., 2022). Flavonoids secreted by legume roots regulate the transcriptional activity of nodulation (*nod*) genes that stimulate rhizobia to produce nodulation factors (NFs), lipochitooligosaccharides, with the backbone of N-acetylglucosamine units, and fatty acids at the non-reducing end. NFs are essential for host-specificity, rhizobial infection, and nodule organogenesis (Dénarié and Cullimore, 1993; Clúa *et al*., 2018; Kidaj *et al*., 2020). Perception of the correct NF structure by compatible receptors in legume root cells initiates early steps of nodulation. These early nodulation events include intracellular calcium oscillations, deformations of root hairs as well as alternations to the root hair cytoskeleton, preparing the host legume plant for symbiotic infection by invading rhizobia (Gage, 2004; Timmers, 2008; Roy *et al*., 2020). Simultaneously, cell divisions in the root cortex and pericycle are reinitiated, leading to the establishment of root nodule primordium with active meristem (Jones *et al*., 2007).

Nodule formation requires two separate, but spatially and temporally highly coordinated processes, namely rhizobial infection of root hairs and nodule organogenesis in the root cortex (Oldroyd and Downie, 2008; Ibáňez *et al*., 2017). Before nodules arise as newly formed functional and nitrogen-fixing root lateral organs, rhizobia must travel from the root surface toward the target cells in the inner root tissue. In the initial stage, rhizobia attach to the growing root hair tips and are trapped in the root hair curls creating enclosed chambers, known as infection pockets (Fournier *et al*., 2015; Rae *et al*., 2021). Within the infection pockets, rhizobia divide and form colonies referred to as infection foci from which infection threads (ITs) entering root hairs are initiated by inverted tip growth. These plant-made tube-like membrane channels are filled with rhizobia, grow down towards the base of infected root hair, and branch out by growing through the root cortex. Eventually, the inward-growing IT and the outward-growing root nodule primordium meet inside the root tissue (Fournier *et al*., 2008; Rashid *et al*., 2015). When ITs reach the nodule primordium, rhizobia are released into the cytoplasm of host cells by endocytosis, become surrounded by plant-derived peribacteroid membrane, and differentiate into bacteroids that are responsible for nitrogen fixation by the activity of nitrogenase enzymatic complex (Terpolilli *et al*., 2012; Poole *et al*., 2018). Since only a specific NFs mixture allows a rhizobial strain to nodulate a particular legume host, mutual compatibility between the two symbionts is essential for establishing a successful symbiotic partnership (Wang *et al*., 2018; Walker *et al*., 2020).

Within a complex signaling network controlling the nodulation process, mitogen-activated protein kinases (MAPKs) become activated early after rhizobial infection (Lopez-Gomez *et al*., 2012). MAPK cascades represent conserved and universal signaling hubs transducing external stimuli into target substrates by a sequential action of three protein kinases, MAPK kinase kinase (MAPKKK), MAPK kinase (MAPKK), and MAPK. Plant MAPKs can be activated by a variety of biotic and abiotic stress stimuli. During signal transduction, MAPKKK reversibly activates its downstream MAPKK, which phosphorylates and activates MAPK by dual phosphorylation of threonine (T) and tyrosine (Y) residues of the T-X-Y motif (Pitzschke, 2015; Xu and Zhang, 2015; Zhang and Zhang, 2022). Activated MAPKs phosphorylate and regulate many diverse substrates such as transcription factors, enzymes, cytoskeletal proteins, or other kinases. Signaling through MAPK modules regulates a broad range of cellular and developmental processes as well as pathogenic or beneficial biotic interactions (Rasmussen *et al*., 2012; Šamajová *et al*., 2013; Smékalová *et al*., 2014; Komis *et al*., 2018; Sun and Zhang, 2022).

Although MAPK-mediated phosphorylation cascades represent an essential component of plant cell signaling, still relatively little is known about MAPKs in legume crops. In alfalfa, stress-induced MAPK (SIMK) is activated by biotic and abiotic stimuli such as fungal elicitors and salt stress, respectively (Munnik *et al*., 1999; Cardinale *et al*., 2000, 2002). Activation analyses and yeast two-hybrid screening identified SIMK kinase (SIMKK) as SIMK specific activator. SIMKK directly activates SIMK in response to salt stress (Kiegerl *et al*., 2000; Cardinale *et al*., 2002) and localization studies at the subcellular level revealed substantial relocation of both SIMKK and SIMK from nuclei to the cytoplasmic spot-like compartments upon salt stress (Ovečka *et al*., 2014). In addition, activated SIMK relocates from nuclei to the tips of growing root hairs and together with the dynamic actin cytoskeleton regulates alfalfa root hair tip growth (Šamaj *et al*., 2002, 2003). Most importantly, we have recently addressed SIMK positive role in alfalfa nodulation and development through its genetic manipulations. For this functional assessment, a transgenic *SIMKK-RNAi* line with a strong downregulation of SIMK production and activity, and a transgenic GFP-SIMK line constitutively overexpressing GFP-tagged and activated SIMK were utilized. SIMK overexpression promoted root hair growth, ITs and nodules clustering, as well as positively affected agronomical traits such as shoot biomass production, suggesting the biotechnological potential of this kinase (Hrbáčková *et al*., 2021). However, the functional and spatiotemporal mode of SIMK participation at early nodulation stages remains unknown.

In this study, live-cell imaging using light-sheet fluorescence microscopy supplemented with quantitative microscopy employing alfalfa-adapted immunolabeling techniques revealed SIMK-specific subcellular localization and activation at early stages of alfalfa − *Sinorhizobium meliloti* symbiotic interaction process. SIMK in root hairs targeted the docking site where *S. meliloti* was attached, entrapped, and internalized. We correlated SIMK subcellular localization patterns in root hairs at *S. meliloti* internalization sites in two contrasting alfalfa genotypes, stably overexpressing GFP-tagged SIMK and downregulating SIMK level by means of *SIMKK* RNAi technology, respectively. SIMK genetic manipulation, the mode of activation, and the localization pattern indicate that the effectiveness of early nodulation steps in alfalfa is modulated by precise spatiotemporal SIMK localization and activation.

## Results

### SIMK distribution in alfalfa growing root hairs

To characterize SIMK localization patterns in growing root hairs of alfalfa control and transgenic plants, whole-mount immunofluorescence analysis after plant fixation was performed. Under control conditions, when alfalfa root hairs were not exposed to *S. meliloti*, a tip-focused pattern of SIMK distribution was observed (Figure 1). Immunodetection revealed mainly apical and sub-apical localization of SIMK in growing root hairs of alfalfa RSY plants (Figure 1A) and plants of transgenic *SIMKK-RNAi* (Figure 1D) and GFP-SIMK (Figure 1G) lines. Moreover, the activated pool of MAPKs in root hairs of RSY (Figure 1B), *SIMKK-RNAi* (Figure 1E) and GFP-SIMK (Figure 1H) plants were spatially detected, showing the same distribution (Figure 1, C, F, I). GFP localization in fixed root hairs of transgenic GFP-SIMK line confirmed the SIMK localization pattern obtained by immunolabeling (Figure 1, G and J). Profiling of fluorescence intensity distribution along individual root hairs was documented by semi-quantitative measurements showing higher accumulation of SIMK and activated MAPKs in the apex and sub-apex of alfalfa root hairs (Figure 1, K-P). In comparison to RSY root hairs (Figure 1, K and L), the displayed profile distribution revealed decreased fluorescence intensity of both SIMK and activated MAPKs in root hair tips of transgenic *SIMKK-RNAi* line (Figure 1, M and N), while the fluorescence intensity of both SIMK and activated MAPKs was increased in root hair tips of transgenic GFP-SIMK line overexpressing GFP-tagged SIMK (Figure 1, O and P). These results demonstrate a considerably decreased presence of activated SIMK in root hair tips of transgenic *SIMKK-RNAi* line compared to RSY, while it was considerably increased in overexpression GFP-SIMK lines. Since root colonization by *S. meliloti* initiates from growing root hairs, the presence of activated SIMK in the root hair tip may be an essential component of initial attachment and invasion steps by rhizobia and could be potentially required for the establishment and efficient formation of early symbiotic structures.

**Figure 1.**
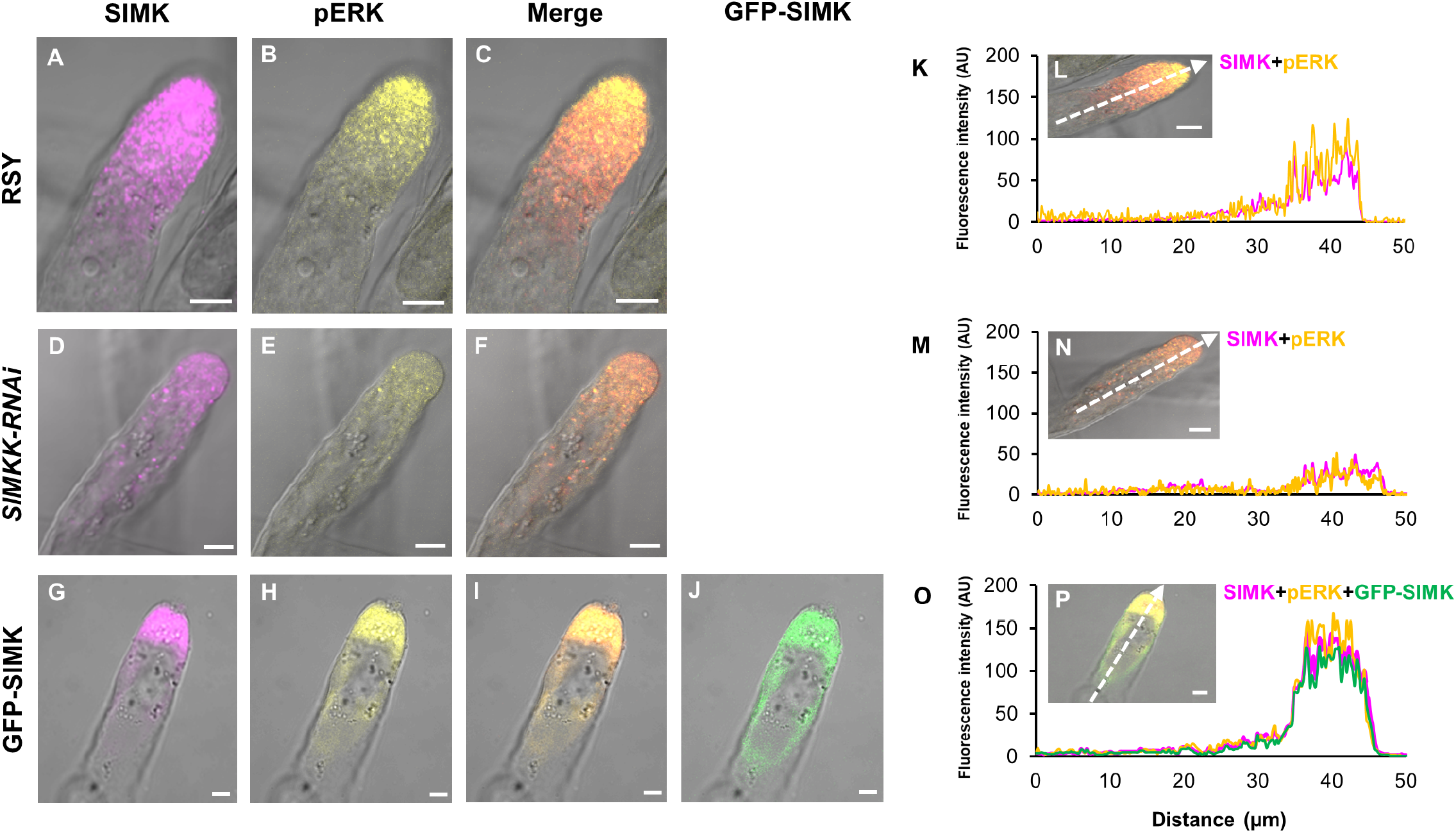
Subcellular immunolocalization of SIMK and activated MAPKs in growing root hairs of alfalfa control and transgenic plants under control conditions. **(A-J)** Whole-mount immunofluorescence localization of SIMK and activated MAPKs in root hairs of alfalfa RSY plants **(A-C)** and plants of transgenic *SIMKK-RNAi* **(D-F)** and GFP-SIMK **(G-J)** lines. SIMK (in magenta) was immunostained with SIMK-specific antibody **(A**,**D**,**G)** and activated MAPKs (pERK, in yellow) using phospho-specific pERK 44/42 antibody **(B**,**E**,**H)**. Overlay of bright-field images with fluorescence channels **(C**,**F**,**I)**, GFP-tagged SIMK (in green) was localized via GFP fluorescence in fixed root hair of transgenic GFP-SIMK line **(J). (K**,**M**,**O)** Fluorescence intensity profiles of SIMK, activated MAPKs, and GFP-tagged SIMK distribution along the measured line shown in **(L**,**N**,**P)**. Scale bar = 5 μm **(A-J; L**,**N**,**P)**.

### The reaction of GFP-SIMK to rhizobia infection

To characterize the GFP-SIMK localization pattern during early nodulation stages *in vitro*, live cell imaging of alfalfa roots stably expressing GFP-tagged SIMK, co-cultivated with mRFP-labeled *S. meliloti* was performed by LSFM 3 to 4 days post inoculation (dpi) (Figure 2A). The mode of interaction was analyzed in the mature part of roots, typically involved in nodulation, where GFP-SIMK is no more accumulated at the tip of root hairs because of terminated tip growth (Figure 2, A and B). GFP-SIMK at this stage is located mainly in nuclei and cytoplasm of root hairs (Hrbáčková *et al*., 2021). Roots inoculated with *S. meliloti* were cultivated on the surface of agar plates, leading to the formation of a dense layer-like biofilm of mRFP-labeled *S. meliloti* associated only with the root and root hairs touching the agar surface. This model allows studying simultaneously and independently root hairs symbiotically interacting with rhizobia, but also root hairs untouched by rhizobia that were exposed to air inside the Petri dish. Both types of root hairs are present on the same root exposed to the same conditions and treatments (Figure 2A). Therefore, 3D rendering of symbiotically infected GFP-SIMK root enabled us to distinguish not only non-interacting alfalfa root hairs from those that interact with rhizobia but also the position of their nuclei with respect to infection (Supplemental Movie S1). In non-growing root hairs that were not in physical contact with rhizobia, nuclei were positioned almost uniformly near the root hair base (Figure 2B; Supplemental Movie S1) while in reactivated root hairs under symbiotic interaction, nuclei were located closer to the site of rhizobia attachment at the tip (Figure 2C; Supplemental Movie S1). Preference for detailed LSFM live-cell imaging was given to the curled root hairs with attached rhizobia (Figure 2D) or rhizobia already enclosed in the root hair curls (Figure 2, E and F), for characterization of SIMK involvement in early infection structures.

**Figure 2.**
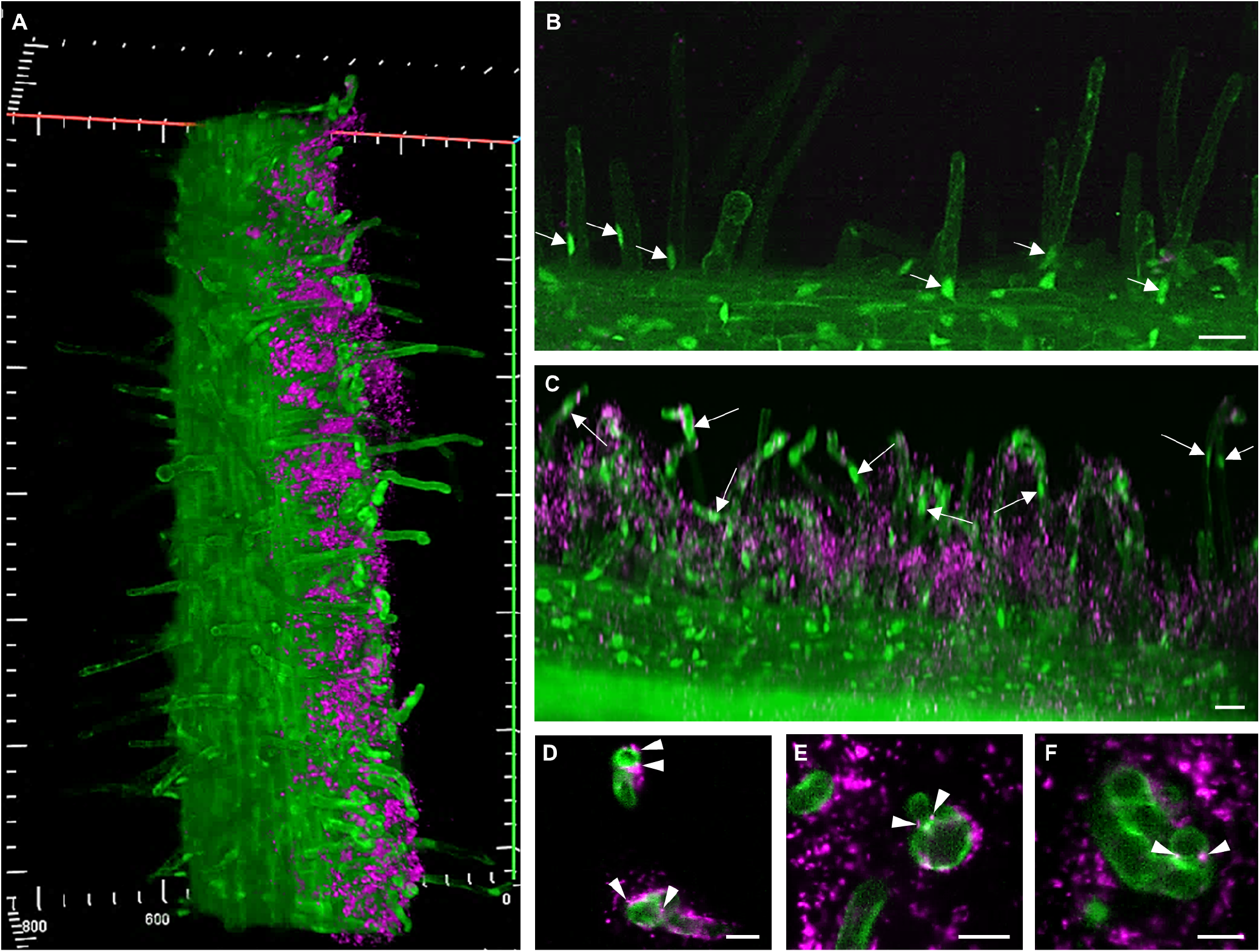
Live cell imaging of early nodulation stages in roots of transgenic GFP-SIMK line at 3 dpi with mRFP-labeled *S. meliloti* using LSFM. **(A)** 3D rendering overview of alfalfa root stably expressing GFP-tagged SIMK (green) co-cultivated with mRFP-labeled rhizobia (magenta). Rhizobia growing on the surface of agar plates are associated with a portion of the root and root hairs that were in contact with the agar plate surface. **(B)** Position of nuclei in root hairs not interacting with rhizobia (arrows). **(C)** Position of nuclei in root hairs interacting with rhizobia (arrows). **(D-F)** Details of root hair infection during rhizobia attachment and internalization (arrowheads). Scale bar = 20 μm **(D-F)**, 40 μm **(B-C)**.

### Rhizobia-induced GFP-SIMK subcellular relocation

To find out whether exposure of alfalfa plants to beneficial rhizobia leads to changes in GFP-SIMK distribution that could be related to the early symbiotic infection, the mean fluorescence intensity of GFP-SIMK was quantitatively evaluated in non-interacting (Figure 3A) and rhizobia-interacting (Figure 3B) root hairs. Under control conditions, GFP-SIMK was present in nuclei and root hair tips of non-interacting root hairs. Nevertheless, a higher amount of GFP-SIMK was detected in nuclei (Figure 3C). During early nodulation stages, GFP-SIMK was localized in nuclei, but substantial accumulation occurred also around rhizobia at specific infection sites (Figure 3B). Upon this early rhizobial infection, the amount of GFP-SIMK in nuclei decreased compared to control conditions (Figure 3 D and E). It seems that GFP-SIMK rather redistributes in root hairs upon rhizobia interaction and accumulates at infection sites where the nodulation process begins (Figure 3, B and D). This finding further suggests the supportive role of SIMK during the early stages of alfalfa nodulation.

**Figure 3.**
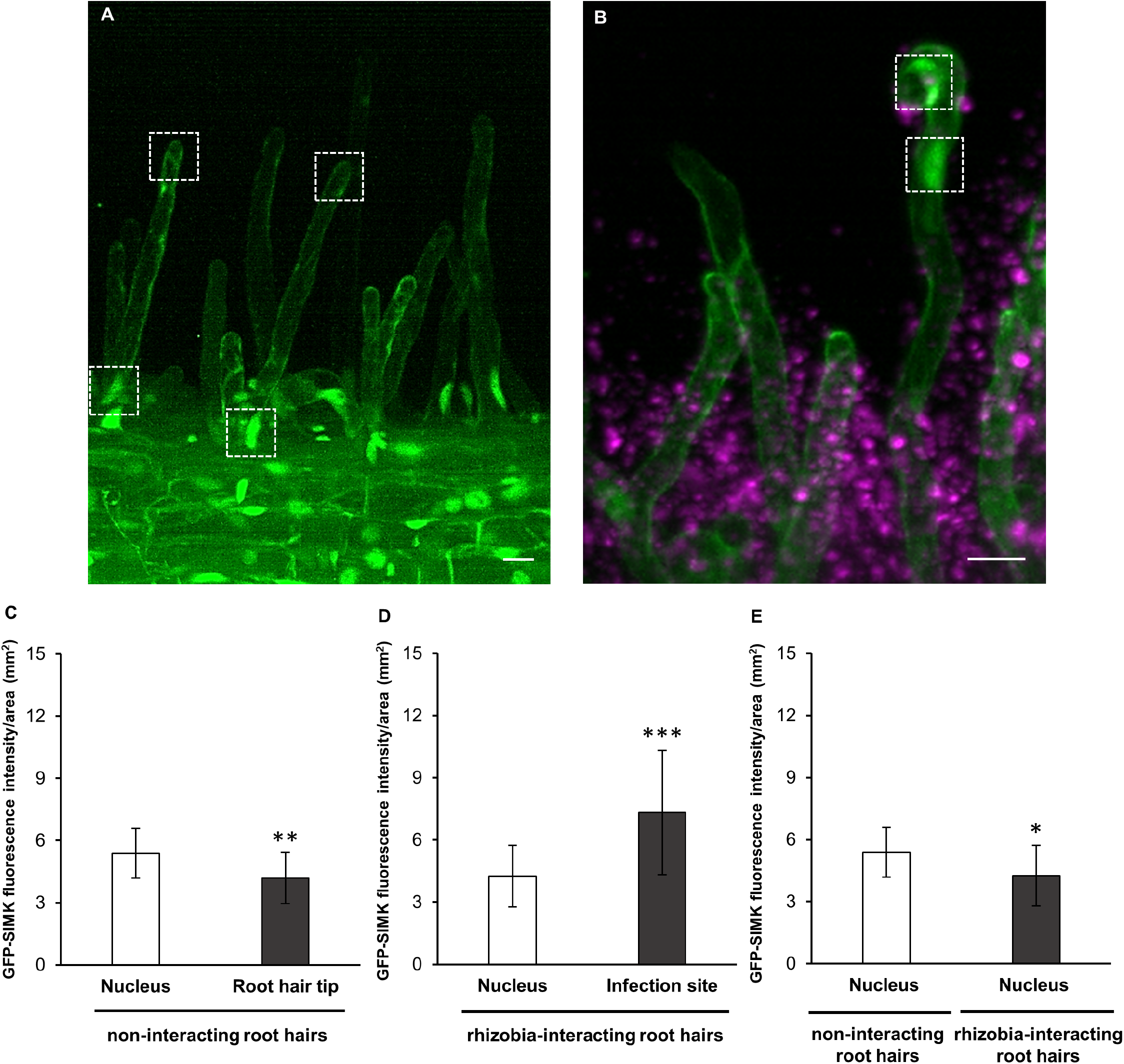
Quantitative analysis of GFP-SIMK fluorescence intensity distribution in nuclei and tips of alfalfa root hairs under control conditions and upon symbiotic interaction with mRFP-labeled *S. meliloti*. **(A-B)** GFP-SIMK distribution in non-interacting root hairs **(A)** and root hairs symbiotically interacting with rhizobia **(B)**. Measurement of GFP-SIMK mean fluorescence intensity was performed in nuclei and root hair tips of non-interacting root hairs **(A**, marked by white dashed boxes) and in nuclei and infection sites of interacting root hairs **(B**, marked by white dashed boxes**). (C-E)** Quantitative evaluation of GFP-SIMK signal intensity in non-interacting root hairs **(C**, N=6) and in interacting root hairs **(D**, N=6), and comparison of GFP-SIMK signal intensity in nuclei of non-interacting and interacting root hairs **(E**, N=6). Statistical differences were calculated in Microsoft Excel using t-test. Error bars show ± standard deviation (SD). Asterisks indicate statistical significance between treatments (*p < 0,05, **p < 0,01, ***p < 0.001). Scale bar = 20 μm **(A-B)**.

### Association of GFP-SIMK with rhizobia infection sites

Detailed live cell imaging of symbiotically-interacting root hairs revealed localization of GFP-SIMK and its association with the position of fluorescently labeled *S. meliloti* at individual early infection stages, beginning from rhizobia attachment (Figure 4A), rhizobia entry into the root hairs (Figure 4, B and C), infection pocket formation (Figure 4D) up to rhizobia complete enclosure inside infection pockets (Figure 4E). The first morphological response to attached rhizobia was root hair curling (Figure 4F). Semi-quantitative evaluation of GFP-SIMK fluorescence intensity distribution showed increased accumulation of GFP-SIMK in the apex of curled root hair, but also at a specific site of rhizobia attachment (Figure 4, F and G). Moreover, orthogonal projections revealed a very close association of GFP-SIMK with rhizobia attached to the root hair at this specific infection site in the X-Z view (Figure 4H; Supplemental Movie S2 at 0:00:14s− 0:00:21s) and the Y-Z view (Figure 4I; Supplemental Movie S3 at 0:00:14s− 0:00:21s). Later upon infection, a cluster of rhizobia was located specifically at the neck of root hair curl where rhizobia will enter the root hair (Figure 4J). Profile measurements revealed an accumulation of GFP-SIMK in the nucleus located close to the infection site, its relocation from the apex, and specific accumulation at the infection site (Figure 4, J and K). Orthogonal projections from the X-Z view (Figure 4L; Supplemental Movie S4 at 0:00:14s− 0:00:18s) and the Y-Z view (Figure 4M; Supplemental Movie S5 at 0:00:16s− 0:00:24s) revealed close association of GFP-SIMK with rhizobia gathered at the root hair curl through which rhizobia internalization typically takes place. Before rhizobia entry into the alfalfa root hairs, a stage of very tight contact between the curled root hair tip and entrapped rhizobia was captured by LSFM (Figure 4N). Profile measurements showed increased fluorescence intensity of GFP-SIMK at the site where rhizobia are in very close contact with the root hair (Figure 4, N and O). Observation of very tight association including the partial overlay was confirmed from orthogonal projections in the X-Z view (Figure 4P; Supplemental Movie S6 at 0:0014s− 0:00:21s) and the Y-Z view (Figure 4Q, Supplemental Movie S7 at 0:00:15s− 0:00:24s). Once were individual rhizobia entrapped inside root hair curl, an infection pocket formation was initiated (Figure 4R). GFP-SIMK was found to be accumulated around rhizobia surrounding them inside root hair curls (Figure 4S) and associated with them as shown from orthogonal projections in the X-Z view (Figure 4T; Supplemental Movie S8 at 0:00:13s− 0:00:25s) and the Y-Z view (Figure 4U; Supplemental Movie S9 at 0:00:14s− 0:00:22s). Later, rhizobia divide and form colonies within infection pockets (Figure 4V) from which ITs are subsequently initiated. GFP-SIMK was strongly accumulated around infection pockets containing rhizobia (Figure 4W) and orthogonal projections in the X-Z view (Figure 4X; Supplemental Movie S10 at 0:00:15s− 0:00:22s) and the Y-Z view (Figure 4Y; Supplemental Movie S11 at 0:00:14s− 0:00:21s) corroborated close GFP-SIMK distribution around infection pockets. Importantly, *in vivo* time-lapse imaging showed that accumulation of GFP-SIMK at the infection site in root hairs during rhizobia attachment (Supplemental Movie S12) and around infection pockets (Supplemental Movie S13) is stable over time, as semi-quantitatively documented profile measurements of GFP-SIMK fluorescence intensity distribution did not fluctuate (Supplemental Movie S14 and S15). Altogether, live cell LSFM imaging showed specific localization and accumulation of GFP-SIMK at infection sites during the early infection stages, which very closely associated with attaching and internalizing rhizobia. Based on the presence of activated SIMK in the growing tips of alfalfa root hairs (Figure 1), it seems that accumulation and activation of SIMK might play an important role in the early stages of alfalfa root hair infection by *S. meliloti*.

**Figure 4.**
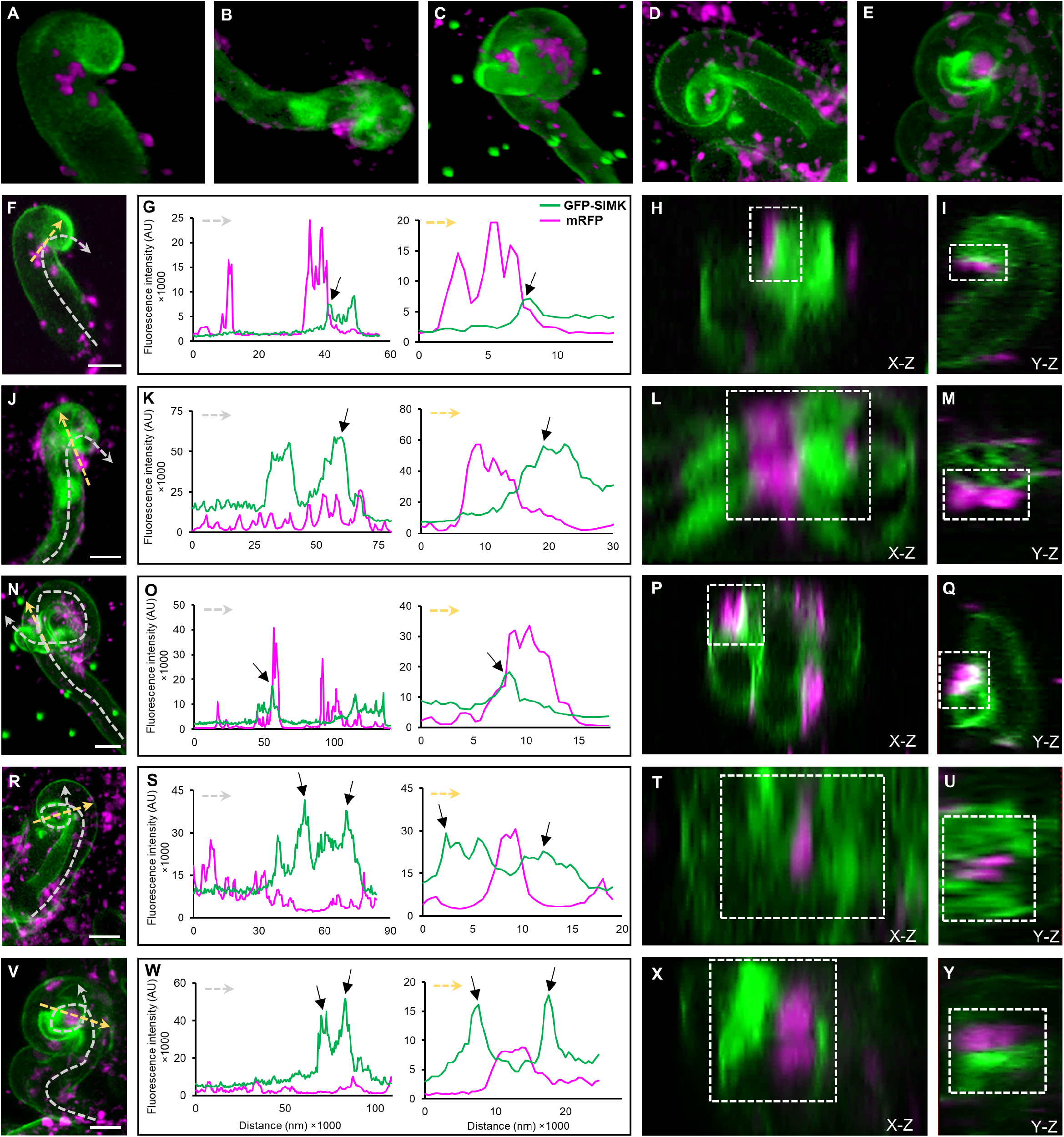
Live cell localization of GFP-SIMK and its association with mRFP-labeled *S. meliloti* during early nodulation stages in root hairs 3 to 4 dpi using LSFM. **(A-E)** Selected root hairs at early sequential infection stages showing the distribution of GFP-SIMK (green) and rhizobia (magenta) during attachment to the root hairs **(A)**, followed by rhizobia internalization **(B, C)**, infection pocket formation **(D)**, and rhizobia enclosure inside infection pockets **(E). (F-Y)** Detailed qualitative and semi-quantitative analysis of GFP-SIMK (green) and rhizobia (magenta) distribution during attachment to the root hairs **(F-I)**, rhizobia internalization **(J-Q)**, infection pocket formation **(R-U)**, and rhizobia enclosure inside infection pockets **(V-Y)**. Semi-quantitative evaluation of GFP-SIMK and mRFP-labeled rhizobia fluorescence distribution **(G**,**K**,**O**,**S**,**W)** along dashed arrows in **(F**,**J**,**N**,**R**,**V)**, indicating GFP-SIMK distribution in symbiotically infected root hairs (grey dashed arrows) and its association with rhizobia at specific infection sites (yellow dashed arrows). Representative images prepared from orthogonal projections in X-Z views **(H**,**L**,**P**,**T**,**X)** and Y-Z views **(I**,**M**,**Q**,**U**,**Y)** show a detailed view of GFP-SIMK accumulation around fluorescently labeled rhizobia (marked with a white dashed boxes). Black arrows in **(G**,**K**,**O**,**S**,**W)** show increased accumulation of GFP-SIMK. Green dots in **(C**,**N)** are fiducial markers. Scale bar = 10 μm **(F**,**J**,**N**,**R**,**V)**.

### SIMK subcellular localization during infection pockets formation

To reveal the subcellular localization pattern of SIMK and activated MAPKs in root hairs of control RSY and transgenic *SIMKK-RNAi* and GFP-SIMK plants 3-7 dpi with *S. meliloti*, and their association with infection pockets, immunofluorescence localization microscopy was employed. Fixed root samples were immunolabeled for SIMK and activated MAPKs using SIMK-specific and phospho-specific antibodies, respectively. The pattern of SIMK and activated MAPK localization was documented in infection pockets, the first symbiotic structure, formed inside root hairs after rhizobial internalization. DAPI, typically used for DNA nuclear staining, effectively stained also *S. meliloti*, which enabled a detailed study of MAPKs association with intracellular compartments enclosing rhizobia during the early stages of the nodulation process. Upon root hair curling, *S. meliloti* was entrapped in alfalfa root hair curls and became completely enclosed inside infection pockets (Figure 5A). At this symbiotic stage, immunostaining revealed SIMK localization close to the plasma membrane and particularly prominent SIMK-specific accumulation around infection pockets in the alfalfa RSY line (Figure 5B). Labeling of activated MAPKs showed the same pattern of localization (Figure 5C), indicating a colocalization with SIMK-specific signal (Figure 5D). This suggests that MAPKs localized around infection pockets were phosphorylated. Moreover, a semi-quantitative evaluation of fluorescence intensity distribution confirmed the close association of both SIMK and activated MAPKs with infection pockets (Figure 5P).

**Figure 5.**
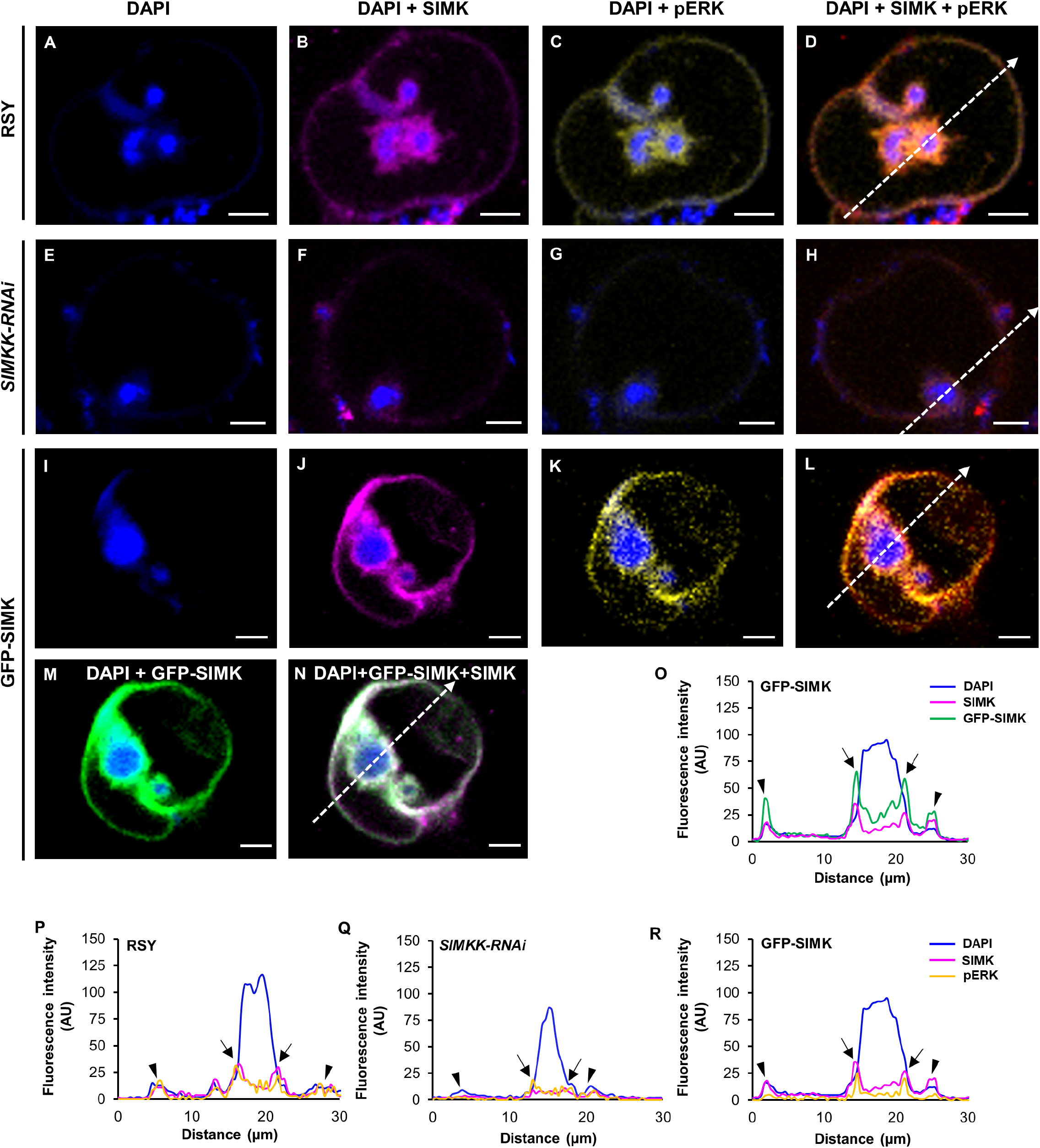
Subcellular immunolocalization of SIMK and activated MAPKs around infection pockets in curled root hairs after inoculation with *S. meliloti*. **(A**,**E**,**I)** Localization of DAPI-stained rhizobia inside infection pocket of RSY **(A)**, *SIMKK-RNAi* **(E)**, and GFP-SIMK **(I)** lines. **(B**,**F**,**J)** SIMK immunostained with SIMK-specific antibody and overlaid with DAPI in RSY **(B)**, *SIMKK-RNAi* **(F)**, and GFP-SIMK **(J)** lines. **(C**,**G**,**K)** Activated MAPKs immunostained with phospho-specific pERK 44/42 antibody and overlaid with DAPI in RSY **(C)**, *SIMKK-RNAi* **(G)**, and GFP-SIMK **(K)** lines. **(D**,**H**,**L)** Overlay of DAPI, SIMK and activated MAPKs in RSY **(D)**, *SIMKK-RNAi* **(H)**, and GFP-SIMK **(L)** lines. **(M**,**N)** GFP-tagged SIMK overlaid with DAPI **(M)** and overlay of GFP-tagged SIMK, SIMK immunostained with SIMK-specific antibody and DAPI in transgenic GFP-SIMK line **(N). (O**,**P**,**Q**,**R)** The fluorescence intensity distribution of SIMK, activated MAPKs, GFP-tagged SIMK, and DAPI was measured along profiles indicated by white dashed arrows in **(D**,**H**,**L**,**N)**. Black arrows indicate the plasma membrane of the infection pocket, black arrowheads indicate the root hair plasma membrane. Scale bar = 5 μm **(A-N)**.

In root hairs of the transgenic *SIMKK-RNAi* line, infection pockets filled with DAPI-stained *S. meliloti* (Figure 5E) were surrounded by a very faint signal of both SIMK (Figure 5F) and activated MAPKs (Figure 5G), showing a similar pattern of localization (Figure 5H). Semi-quantitative profile measurements revealed an association of SIMK and activated MAPKs with infection pockets. However, compared to the alfalfa RSY plants (Figure 5P), the fluorescence intensity of SIMK and activated MAPKs was substantially decreased in the transgenic *SIMKK-RNAi* line (Figure 5Q).

Similarly, inside root hairs of transgenic GFP-SIMK line, when *S. meliloti* became fully entrapped inside infection pockets (Figure 5I), immunodetection with SIMK-specific antibody revealed substantial accumulation mainly around infection pockets, but prominent SIMK-specific signal was also detected at the plasma membrane of curled root hairs (Figure 5J). Activated MAPKs showed a similar subcellular localization pattern around infection pockets and at the plasma membrane of curled root hairs (Figure 5K), leading to a high degree of colocalization with SIMK-specific signal (Figure 5L). SIMK localization pattern obtained by immunostaining with SIMK-specific antibody was independently confirmed by localization of GFP-tagged SIMK in curled root hairs of the GFP-SIMK line (Figure 5M). Also, GFP-tagged SIMK showed a high degree of colocalization with SIMK signal obtained by immunolabeling with SIMK-specific antibody (Figure 5N), and semi-quantitative evaluation of SIMK and activated MAPKs fluorescence intensity distribution clearly revealed its close association with infection pockets (Figure 5, O and R).

In addition, quantitative comparative analysis of mean fluorescence intensity around infection pockets revealed, in comparison to control RSY (Figure 6, A and G), significantly higher levels of SIMK in GFP-SIMK plants (Figure 6, C and G) and significantly lower in transgenic *SIMKK-RNAi* line (Figure 6, B and G). Also, a lower level of activated MAPKs was found around infection pockets inside root hairs of the transgenic *SIMKK-RNAi* line (Figure 6, E and H), while no significant difference was observed in the transgenic GFP-SIMK line compared to alfalfa RSY plants (Figure 6, D, F and H).

**Figure 6.**
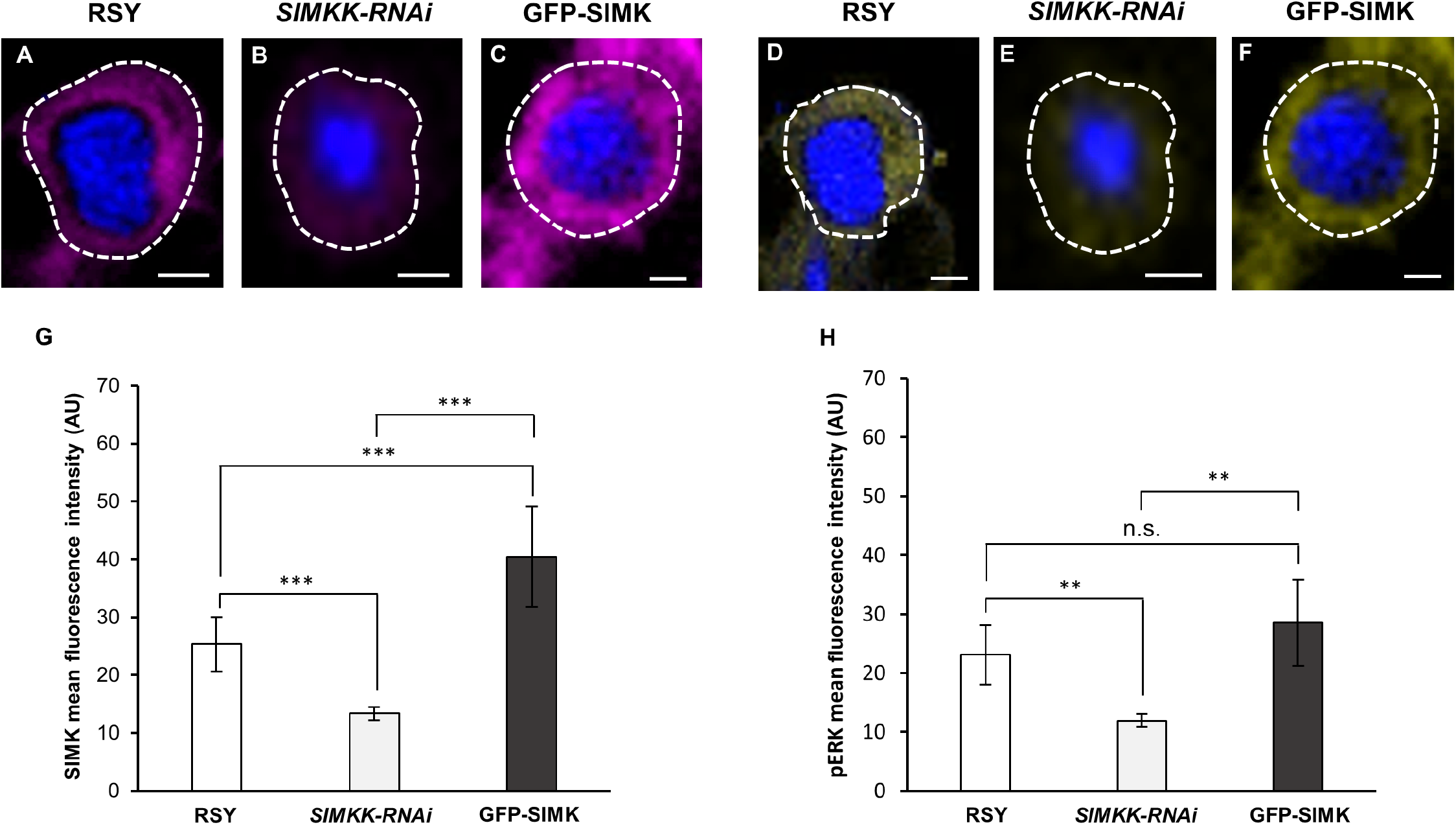
Quantitative analysis of SIMK and phosphorylated MAPKs fluorescence intensity distribution around infection pockets in curled root hairs after inoculation with *S. meliloti*. **(A-F)** Immunolocalization of SIMK **(A-C)** and phosphorylated MAPKs **(D-F)** in infection pockets of RSY **(A**,**D**; N=10 for SIMK, N=7 for pERK), *SIMKK-RNAi* **(B**,**E**; N=10 for SIMK, N=7 for pERK), and GFP-SIMK **(C**,**F**; N=9 for SIMK, N=7 for pERK). White dashed lines in **(A-F)** indicate defined ROIs in which the mean fluorescence intensity was measured. **(G**,**H)** Quantitative evaluation of SIMK **(G)** and activated MAPKs **(H)** signal intensity in transgenic *SIMKK-RNAi* and GFP-SIMK lines compared to RSY plants. Statistical differences were calculated in Microsoft Excel using t-test. Error bars show ± standard deviation (SD). Asterisks indicate statistical significance between treatments (**p < 0,01, ***p < 0.001, n.s. no statistical significance). Scale bar = 2 μm **(A-F)**.

Quantitative determination of the colocalization rate between signals of SIMK and activated MAPKs, expressed by Pearson’s correlation coefficient, revealed the highest value around infection pockets in the transgenic GFP-SIMK line, and the lowest in transgenic *SIMKK-RNAi* line (Supplemental Figure S1).

This analysis clearly showed SIMK-specific accumulation and activation around infection pockets containing entrapped *S. meliloti* in alfalfa root hairs. The level of active SIMK accumulation was strongly associated with the SIMK expression level. It was substantial in root hairs of the transgenic GFP-SIMK line, while the lowest presence of active SIMK was detected around infection pockets in the transgenic *SIMKK-RNAi* line causing strong downregulation of both *SIMKK* and *SIMK* (Hrbáčková *et al*., 2021). Since infection pockets represent the site of *S. meliloti* entry and ITs initiation, these results indicate that active SIMK accumulated at this specific location might be required for efficient ITs formation.

### SIMK subcellular localization during ITs formation

Complete enclosure of rhizobia inside infection pockets is followed by an invagination of the host cell plasma membrane and initiation of tunnel-like ITs. Therefore, the pattern of SIMK and activated MAPKs subcellular localization was characterized by immunolabeling during ITs formation and propagation through root hairs. Inside root hairs of the alfalfa RSY line, ITs were easily detectable owing to DAPI-stained *S. meliloti* (Figure 7A). Immunostaining revealed SIMK-specific signals surrounding growing ITs (Figure 7B). Activated MAPKs immunolabeled with anti-phospho-p44/42 antibody showed the same subcellular distribution (Figure 7C) leading to a high degree of colocalization with the SIMK signal (Figure 7D). This colocalization pattern suggests that SIMK located around ITs was phosphorylated. Semi-quantitative analysis of the fluorescence intensity distribution revealed a close association of both SIMK and activated MAPKs with the surface of infection threads (Figure 7O).

**Figure 7.**
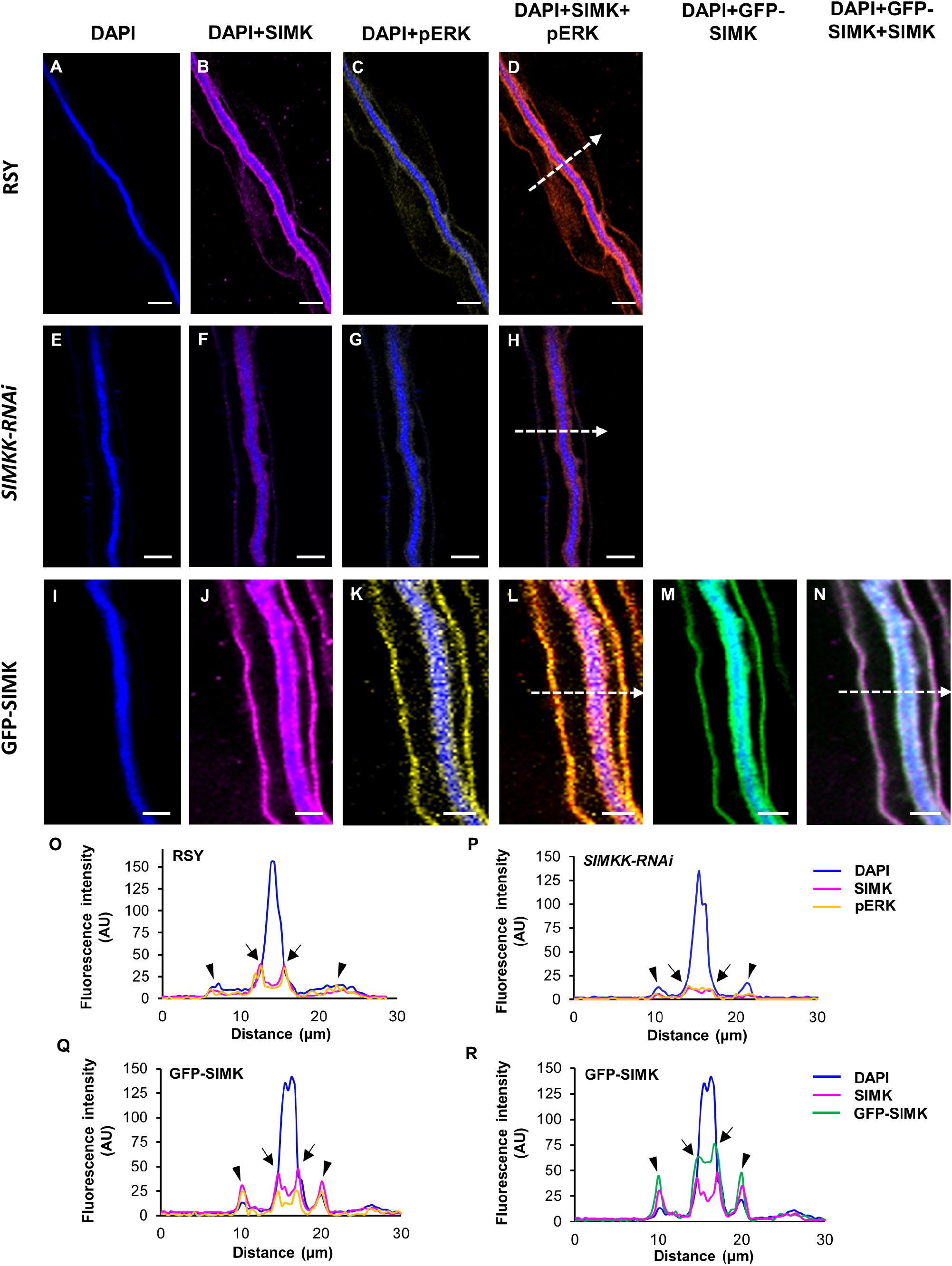
Subcellular immunolocalization of SIMK and activated MAPKs around ITs in root hairs induced after inoculation with *S. meliloti*. **(A**,**E**,**I)** Localization of DAPI-stained rhizobia inside ITs of RSY **(A)**, *SIMKK-RNAi* **(E)**, and GFP-SIMK **(I)** lines. **(B**,**F**,**J)** SIMK immunostained with SIMK-specific antibody and overlaid with DAPI in RSY **(B)**, *SIMKK-RNAi* **(F)**, and GFP-SIMK **(J)** lines. **(C**,**G**,**K)** Activated MAPKs immunostained with phospho-specific pERK 44/42 antibody and overlaid with DAPI in RSY **(C)**, *SIMKK-RNAi* **(G)**, and GFP-SIMK **(K)** lines. **(D**,**H**,**L)** Overlay of DAPI, SIMK and activated MAPKs in RSY **(D)**, *SIMKK-RNAi* **(H)**, and GFP-SIMK **(L)** plants. **(M**,**N)** GFP-tagged SIMK overlaid with DAPI **(M)** and overlay of GFP-tagged SIMK, SIMK immunostained with SIMK-specific antibody and DAPI in transgenic GFP-SIMK line **(N). (O**,**P**,**Q**,**R)** The fluorescence intensity distribution of SIMK, activated MAPKs, GFP-tagged SIMK, and DAPI was measured along profiles indicated by white dashed arrows in **(D**,**H**,**L**,**N)**. Black arrows indicate the plasma membrane of IT, black arrowheads indicate the plasma membrane of root hair. Scale bar = 5 μm **(A-N)**.

In the transgenic *SIMKK-RNAi* line, ITs filled with *S. meliloti* (Figure 7E) were similarly decorated by MAPKs, but showed a very weak signal of both SIMK (Figure 7F) and activated MAPKs (Figure 7G). Nevertheless, the distribution pattern indicated their subcellular colocalization (Figure 7H). Profiling of SIMK and activated MAPKs fluorescence intensity distribution revealed their association with ITs, but substantially decreased (Figure 7P).

In the case of ITs in the transgenic GFP-SIMK line (Figure 7I), immunostaining with SIMK-specific antibody revealed a strong accumulation of SIMK not only along ITs, but also at the plasma membrane of root hairs (Figure 7J). Signal specific for activated MAPKs showed the same pattern of subcellular localization (Figure 7K) and colocalized with SIMK signal (Figure 7L). Observation of GFP-tagged SIMK along ITs (Figure 7M) confirmed the localization pattern obtained by immunolabeling with SIMK-specific antibody and colocalized with SIMK signal (Figure 7N). Semi-quantitative evaluation of fluorescence intensity along indicated profiles (Figure 7, L and N) confirmed enhanced and close association of SIMK and activated MAPKs with ITs and plasma membrane of root hairs (Figure 7, Q and R).

Moreover, the amount of SIMK (Figure 8, A-C) and activated MAPKs (Figure 8, D-F) determined by quantitative analysis of mean fluorescence intensity around ITs was markedly lower in plants of transgenic *SIMKK-RNAi* line in comparison to alfalfa RSY and GFP-SIMK plants (Figure 8, G and H), while the amount of activated MAPKs in transgenic GFP-SIMK line was similar to RSY plants (Figure 8H).

**Figure 8.**
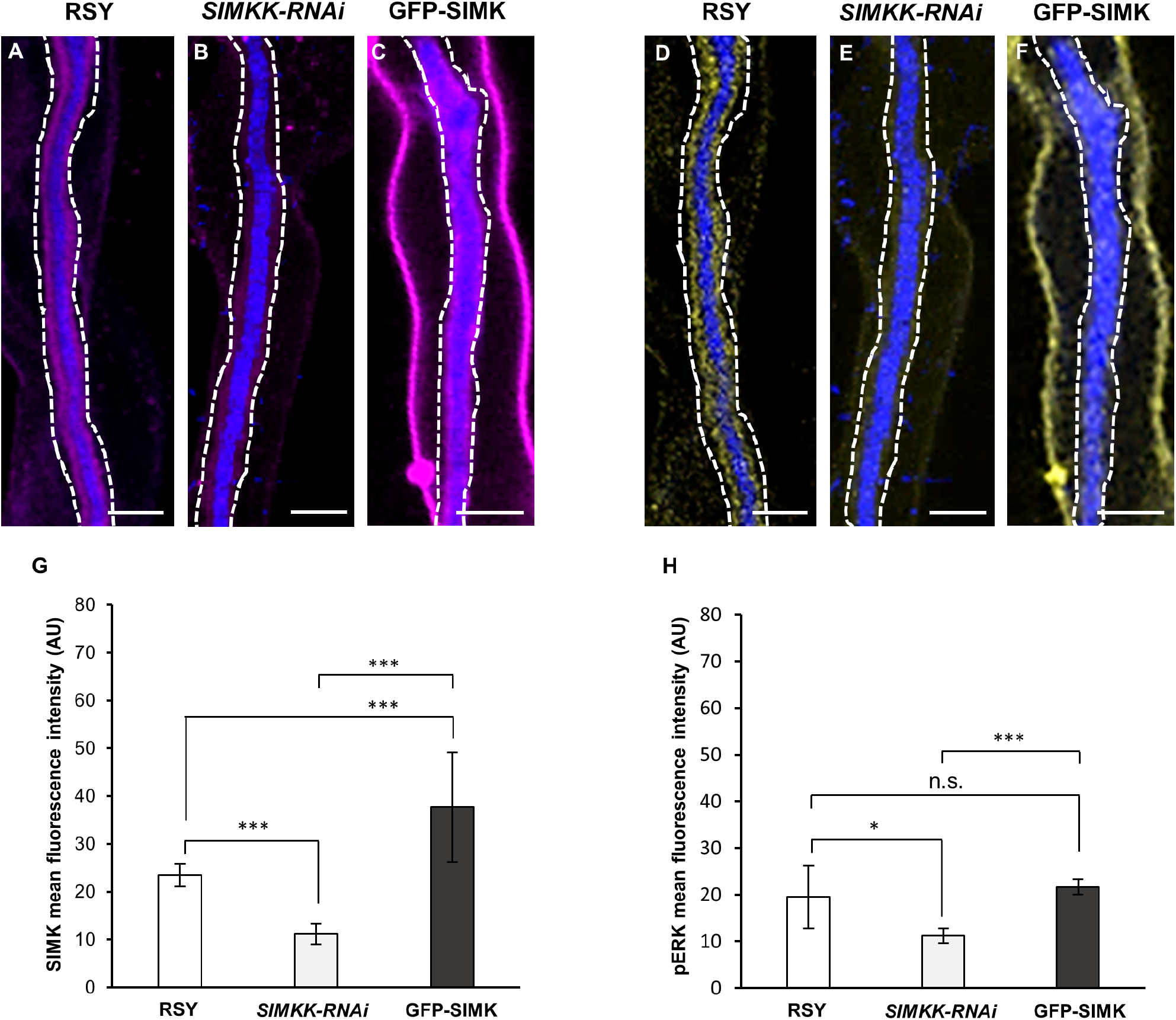
Quantitative analysis of SIMK and phosphorylated MAPKs fluorescence intensity distribution around ITs in root hairs after inoculation with *S. meliloti*. **(A-F)** Immunolocalization of SIMK **(A-C)** and phosphorylated MAPKs **(D-F)** in ITs of RSY **(A**,**D**; N=8 for SIMK, N=8 for pERK), *SIMKK-RNAi* **(B**,**E**; N=6 for SIMK, N=7 for pERK), and GFP-SIMK **(C**,**F**; N=8 for SIMK, N=8 for pERK). White dashed lines in **(A-F)** indicate defined ROIs in which the mean fluorescence intensity was measured. **(G**,**H)** Quantitative evaluation of SIMK **(G)** and activated MAPKs **(H)** signal intensity in transgenic *SIMKK-RNAi* and GFP-SIMK lines compared to RSY plants. Statistical differences were calculated in Microsoft Excel using t-test. Error bars show ± standard deviation (SD). Asterisks indicate statistical significance between treatments (*p < 0,05, ***p < 0.001, n.s. no statistical significance). Scale bar = 5 μm **(A-F)**.

The colocalization rate between SIMK and activated MAPKs was quantitatively determined by Pearson’s correlation coefficient, revealing that the overall colocalization rate between SIMK and activated MAPKs was significantly higher along ITs in the transgenic GFP-SIMK line and RSY plants compared to the transgenic *SIMKK-RNAi* plants (Supplemental Figure S2).

Immunolocalization together with semi-quantitative and quantitative colocalization analyses clearly revealed the presence of SIMK-specific signal along ITs in alfalfa root hairs. Increased accumulation of active SIMK along ITs was observed in the transgenic GFP-SIMK line, while the lowest one was detected in the *SIMKK-RNAi* line. All these data indicate that active SIMK might be involved during ITs formation and its growth toward the place of root nodule primordia initiation. Therefore, SIMK supportive role in the propagation of rhizobia-filled Its through plant root hairs and cortex tissues might play an important role in the regulation and effectiveness of rhizobia delivery to the nodule primordium and subsequent nodule formation.

Moreover, we performed a visualization of plasma membranes using a fixable FM4-64FX, allowing precise observation of SIMK subcellular localization with regard to membranes of early symbiotic structures. Whole-mount immunofluorescence co-labeling in RSY revealed the presence of SIMK close to the membranous surface of infection pockets (Figure 9, A and C) and ITs (Figure 9, B and D). A lower amount of SIMK was found on membranes of infection pockets (Figure 9, E and G) and ITs (Figure 9, F and H) in the *SIMKK-RNAi* line, while a substantially increased amount of SIMK was accumulated on the surface of infection pockets (Figure 9, I and K) and ITs (Figure 9, J and L) in GFP-SIMK line. In quantitative terms, Pearson’s correlation coefficient showed the highest colocalization rate between SIMK and FM4-64FX-labeled membranes in the GFP-SIMK line, while the degree of colocalization was considerably decreased in the *SIMKK-RNAi* line (Figure 9M). The data suggest a close association and interaction of SIMK with membranes of infection pockets and ITs during early nodulation stages.

**Figure 9.**
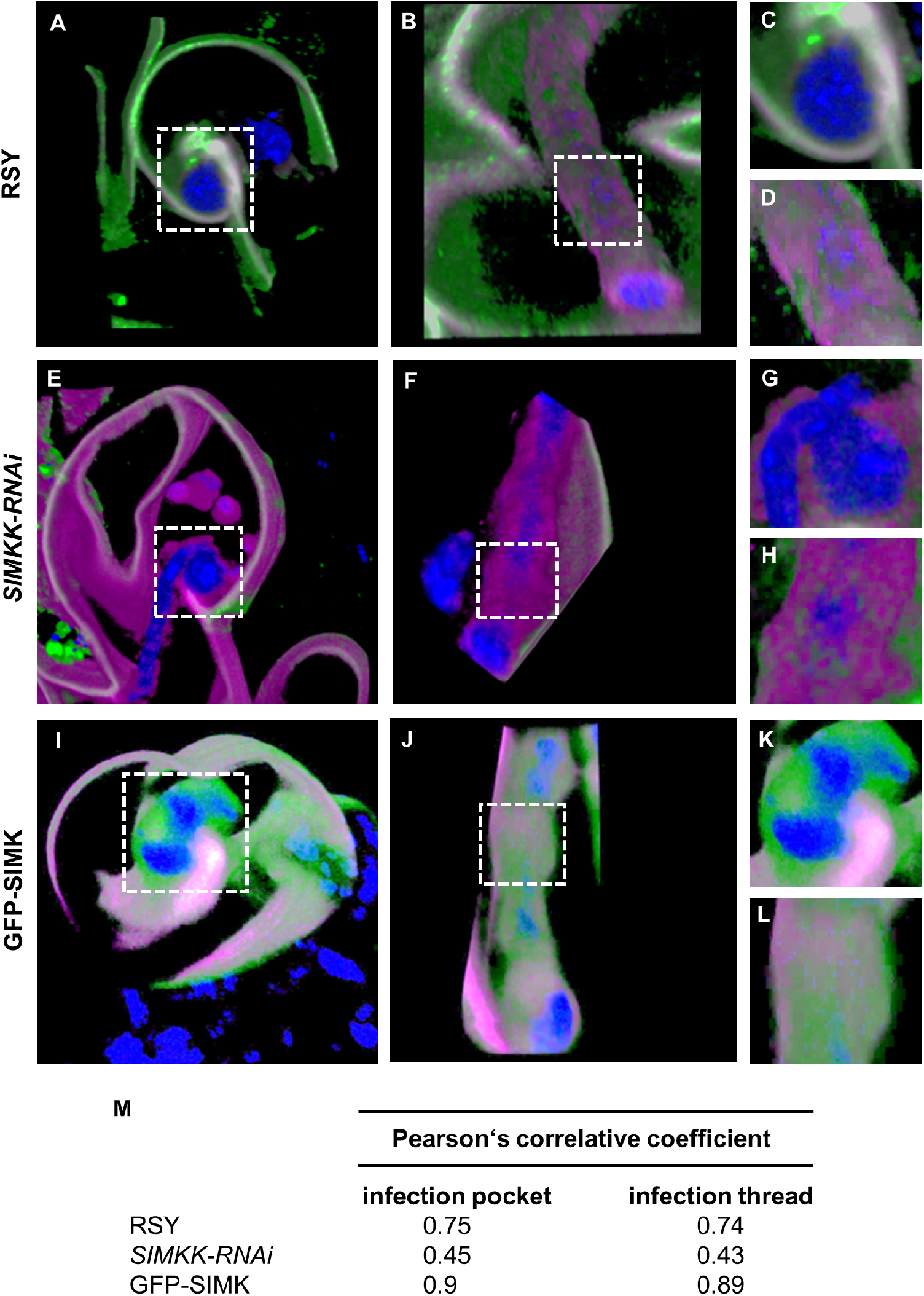
Volume 3D rendering of rhizobia-containing early symbiotic structures with immunolabeled SIMK and membranes visualized using FM4-64FX in root hairs after inoculation with *S. meliloti*. **(A-D)** RSY root hairs. **(E-H)** Root hairs of *SIMKK-RNAi* line. **(I-L)** Root hairs of GFP-SIMK line. Subcellular localization of SIMK with membranes of infection pockets **(A**,**C**,**E**,**G**,**I**,**K)** and ITs **(B**,**D**,**F**,**HJ**,**L)**. Overlay of membranes (in magenta), SIMK (in green) and DAPI-stained rhizobia (in blue). **(M)** Averaged Pearson’s correlative coefficients of colocalization analysis between SIMK and FM4-64FX-stained membranes around infection pockets and ITs. Details of infection pockets and ITs shown in **(C, D, G, H, K, L)** are marked with a white dashed boxes in **(A**,**B**,**E**,**F**,**I**,**J)**.

### Involvement of active SIMK in ITs formation

To correlate the presence of active SIMK at infection pockets and ITs with nodule formation, the efficiency of root hair infection by *S. meliloti* was examined. The number of ITs per the whole root system length was determined in alfalfa RSY and transgenic *SIMKK-RNAi* and GFP-SIMK plants at 10 dpi with *S. meliloti*, when actively growing ITs should already reach the root cortex. No significant difference was observed in the averaged root system length among the three respective lines at 10 dpi with *S. meliloti* (Supplemental Figure S3A). However, the transgenic *SIMKK-RNAi* line, having strongly downregulated both *SIMKK* and *SIMK* (Hrbáčková *et al*., 2021), showed the lowest amount of active SIMK around infection pockets (Figure 6, E and H, Figure 9, E,G,M), formed significantly fewer ITs compared to RSY and transgenic GFP-SIMK plants, while the transgenic GFP-SIMK line produced a similar number of ITs as RSY plants (Supplemental Figure S3B).

## Discussion

Leguminous plants are immensely important to the ecosystem and sustainable agriculture worldwide. Part of their success lies in mutualistic partnership with beneficial nitrogen-fixing bacteria that can convert atmospheric dinitrogen into bioavailable ammonium inside functional root nodules, which helps them to manage nitrogen shortage and facilitate nutrient uptake (Brundrett, 2002; Bisseling and Geurts, 2020). Alfalfa has become a high-quality forage crop with high biological and agronomical potential, especially for its widespread production, ecological adaptability, high nutrition value, and ability to improve nitrogen-limited soils (Radović *et al*., 2009). Early steps of nodulation and subsequent nodule development depend on a molecular dialogue between the host legume and rhizobia, including the exchange of signals and activation of protein-phosphorylation-mediated signal transduction cascades (Shaw and Long, 2003; Grimsrud *et al*., 2010). Specifically in plants, developmental and cellular processes are regulated by MAPK-mediated phosphorylation cascades, and the activity of various protein kinases was shown to be also involved in symbiotic interactions and nodule formation (Grimsrud *et al*., 2010; Komis *et al*., 2018; Roy *et al*., 2020). Although symbiotic nitrogen fixation is extensively studied in model legume species, such as *Medicago truncatula* and *Lotus japonicus*, little is known about the regulation of symbiotic interaction and the possible involvement of MAPK signaling in alfalfa nodulation. Here, we characterized the subcellular localization and activation pattern of SIMK involved in the early stages of alfalfa interaction with *Sinorhizobium meliloti* using genetically engineered transgenic lines.

Despite the remarkable progress in the understanding of MAPK regulation in plant development and immunity, their involvement in various steps of nodule development is still not known. In *L. japonicus*, a MAPKK SIP2 was identified to interact with a symbiosis receptor-like kinase (SymRK), having an essential role in early symbiotic signaling (Chen *et al*., 2012). Yin *et al*. (2019) identified LjMPK6 as the phosphorylation target of SIP2 and showed that the SymRK-SIP2-LjMPK6 signaling module is required for nodule organogenesis and formation in *L. japonicus*. In addition, a recent study demonstrated that LjPP2C, a type 2C protein phosphatase, fine-tunes nodule development in *L. japonicus* via dephosphorylating LjMPK6 (Yan *et al*., 2020). In *M. truncatula*, MtMAPKK4 shows a high sequence identity to MsSIMKK and LjSIP2. Downstream interacting partners of MtMAPKK4 are MtMAPK3 and MtMAPK6, while the MtMAPKK4-MtMAPK3/6 pathway is involved in nodule formation and also with *M. truncatula* general growth and development (Chen *et al*., 2017). Another MAPKK from *M. truncatula*, MtMAPKK5, directly activates MtMAPK3 and MtMAPK6, and the stress signaling-mediated MtMAPKK5-MtMAPK3/6 module negatively affects root nodulation (Ryu *et al*., 2017). In alfalfa, SIMKK is a specific upstream activator of SIMK under salt stress (Kiegerl *et al*., 2000) and both SIMKK and SIMK relocate from the nucleus to cytoplasm under salt stress (Ovečka *et al*., 2014). SIMK overexpression leads to the development of longer root hairs and promoted ITs and nodule clustering (Hrbáčková *et al*., 2021). In contrast, SIMK downregulation was accompanied by the formation of shorter root hairs and few ITs and nodules. Moreover, SIMK overexpression promoted shoot biomass production, and leaf and petiole development. However, a detailed study of SIMK subcellular localization and activation pattern clarifying the spatial and temporal model of SIMK involvement in alfalfa early nodulation stages remained unclear.

Possible SIMK involvement in alfalfa nodulation can be anticipated from its subcellular localization and activation during early symbiotic stages. Crucial is the very tight association of activated SIMK with infection pockets and ITs. We have recently established light-sheet fluorescence microscopy for the spatiotemporal imaging of plant development at subcellular, cellular, tissue and organ levels under controlled environmental conditions (Ovečka *et al*., 2018). It was utilized in alfalfa for the characterization of root development (Vyplelová *et al*., 2018) and the involvement of actin cytoskeleton in the interaction with *S. meliloti* (Ovečka *et al*., 2022). Live-cell imaging using LSFM clearly showed relocation of SIMK from root hair tips to the *S. meliloti* docking site and further close association with sites of rhizobia internalization. We developed also reliable immunolocalization protocols for whole-mount immunolabeling of root samples of *M. sativa*, achieving high signal efficiency and superb sample stability (Tichá *et al*., 2020). Employing these immunolabeling methods explicitly adapted for alfalfa plantlets originating from somatic embryos, we show here subcellular localization patterns of SIMK during the early stages of the nodulation process in alfalfa. Moreover, immunolocalization of phosphorylated MAPKs enabled us to check out whether SIMK is activated or not at the subcellular level. Under control conditions, alfalfa RSY plants and plants of transgenic *SIMKK-RNAi* and GFP-SIMK lines showed a tip-focused pattern of activated SIMK localization in growing root hairs in agreement with previously published data on SIMK localization from both live-cell imaging (Hrbáčková *et al*., 2021) and immunofluorescence microscopy (Šamaj *et al*., 2002). However, decreased accumulation of activated SIMK in growing root hair tips was observed in the transgenic *SIMKK-RNAi* line, consistent with Bekešová *et al*. (2015) showing overall decreased accumulation of phosphorylated SIMK in *SIMKK-RNAi* lines. Although rhizobia can use different routes to invade plant roots, entrance via root hairs is probably the best understood and could be found in legumes such as alfalfa, soybean, pea, bean and vetch (Sprent and James, 2007; Sprent *et al*., 2008; Ibáňez *et al*., 2017). Since root hairs make the first contact with symbiotic rhizobia, active SIMK in root hairs may play an important role in the alfalfa’s early interaction with *S. meliloti* during and after rhizobia attachment. Activation of signaling pathways in the epidermal cells leads to localized inhibition of the tip growth of root hairs and induces its physical curling around attached rhizobia. It is followed by the formation of infection pockets and infection threads, structures essential for rhizobia internalization and delivery toward the target cells in newly forming nodules (Brewin, 2004; Gage 2004). In contrast to the overexpressor GFP-SIMK line, where activated SIMK was strongly accumulated around infection pockets and ITs, the transgenic *SIMKK-RNAi* line showed much-decreased accumulation. Indeed, the number of formed ITs was significantly lower in the transgenic *SIMKK-RNAi* line, indicating the importance of activated SIMK in infection pockets, which is further required for proper ITs formation. Therefore, SIMK downregulation negatively affects nodule formation, while SIMK overexpression enhances infection pockets and ITs formation.

Conclusively, we show that active SIMK is associated with *S. meliloti* internalization sites in root hairs and with ITs delivering *S. meliloti* to internal root tissues. SIMK downregulation negatively affects infection pockets and ITs formation. The subcellular immunolocalization pattern supported by the localization pattern of GFP-SIMK in living cells thus clearly demonstrates that active SIMK might be a key player responsible for fine-tuning of the nodulation process in alfalfa. SIMK, therefore, represents a potentially new regulatory protein required for the establishment of efficient symbiotic interaction in alfalfa.

## Materials and methods

### Plant material

Alfalfa wild-type plants of cv. Regen-SY (RSY) carrying either *35S::GFP:SIMK* construct (GFP-SIMK fusion protein; Hrbáčková *et al*., 2021) or *SIMKK-RNAi* in pHellsgate12 plasmid driven under *35S* promoter (obtained from CSIRO Plant Industry, Australia) were obtained by regeneration *in vitro* through somatic embryogenesis from leaf explants as previously described (Samac and Austin-Phillips, 2006; Bekešová *et al*., 2015; Hrbáčková *et al*., 2021). Regenerated alfalfa plants RSY, transgenic *SIMKK-RNAi* line (showing strong downregulation of *SIMKK* and *SIMK* transcripts and SIMK protein), and transgenic GFP-SIMK line with upregulated *SIMK* transcript and enhanced SIMK activity (Hrbáčková *et al*., 2021) were transferred to nitrogen-free Fåhreus medium (FAH-N_2_; Fåhreus, 1957) for inoculation with rhizobia.

### Plant inoculation with S. meliloti

Regenerated plantlets of transgenic GFP-SIMK line approximately 1.5 cm long and growing on a FAH-N_2_ medium containing 13 g/L of micro agar were inoculated with *S. meliloti* (strain Sm2011) producing mRFP with OD_600_= 0.5 (2.50e+008 cell/ml) and used for live-cell imaging 3 to 4 dpi. For quantitative analyses, 18 days old plants of alfalfa RSY and transgenic *SIMKK-RNAi* and GFP-SIMK lines originating from somatic embryos and growing on the FAH-N_2_ medium were inoculated with *S. meliloti* wild-type (strain Sm2011) with OD_600_= 0.5. In total, 2 ml of rhizobial suspension was applied to the root system directly on plates, followed by vertical cultivation of inoculated plants in an environmental chamber at 21°C, 70% humidity, and 16h/8h light/dark cycle with covered root systems.

### Sample preparation for live-cell imaging and light-sheet fluorescence microscopy (LSFM)

Plantlets of transgenic GFP-SIMK line were used for live cell imaging to observe *in vivo* localization and dynamics of GFP-tagged SIMK during the early stages of nodulation. Samples for LSFM imaging were prepared according to Ovečka *et al*., 2015. Fluorinated ethylene propylene (FEP) tube with an inner diameter of 4.2 mm and outer diameter of 4.6 mm was connected to the glass capillary (inner diameter of 2.15 mm and outer diameter of 4.0 mm) using the hot glue gun. Inoculated plantlet was gently inserted into the FEP tube with tweezers and medium (FAH-N_2_ medium pH 6.5) with 1% (w/v) low melting point agarose (Sigma Aldrich) containing fiducial markers (fluorescent beats of 1 μm in diameter) was slowly added from the bottom into the FEP tube. Under these conditions, the plant root was embedded in the solidified block of the culture medium inside the FEP tube while the green upper part of the plant was exposed to air. The glass capillary connected to the FEP tube containing the embedded sample was fixed into the metal holder and directly placed into a pre-tempered (22°C) LSFM observation chamber filled with a liquid FAH-N_2_ medium. After sample stabilization, imaging was performed using the light-sheet Z.1 fluorescence microscope (Carl Zeiss, Germany) equipped with Plan-Apochromat 10×/0.5 NA detection objective and two LSFM 10×/0.2 NA illumination objectives (Carl Zeiss, Germany). Rhizobia-infected roots were imaged using dual-side light-sheet illumination with excitation laser lines 488 nm for GFP (beam splitter LP 560 and emission filter BP 505-545) and 561 nm for RFP (beam splitter LP 560 and emission filter BP 575-615). Images were acquired with the PCO.Edge sCMOS cameras (PCO AG, Germany) with an exposure time of 50 ms and an imaging frequency of every 2 min in Z-stack mode for 80 and 120 min. The scaling of acquired images in x, y, and z dimensions was 0.466 μm × 0.466 μm × 1.497 μm, and light-sheet thickness was set to the optimal value.

### Fixation of alfalfa root samples

For whole-mount immunofluorescence labeling, approximately 1.5 cm long root segments were excised from roots of alfalfa RSY, *SIMKK-RNAi* and GFP-SIMK plants co-cultivated with *S. meliloti* and fixed in freshly prepared fixative solution (Tichá *et al*., 2020). Sampling was done at 3-7 dpi with *S. meliloti* when infection pockets and growing ITs were clearly detectable inside rhizobia-infected root hairs after microscopic observation.

### Immunolabeling of SIMK and activated MAPKs in symbiotically-infected root hairs

SIMK subcellular localization at early symbiotic stages was performed by immunofluorescence labeling on fixed root samples of alfalfa RSY, *SIMKK-RNAi* and GFP-SIMK lines co-cultivated with *S. meliloti* using a SIMK-specific antibody. To check out the activation state of SIMK in analyzed early stages of nodulation, an activated pool of MAPKs was immunodetected using a phospho-specific antibody (anti-phospho-p44/42, Cell Signaling, Netherlands). Root samples were simultaneously double-immunolabeled with rabbit anti-AtMPK6 (SIMK-specific) primary antibody (Sigma, Life Science, USA) at 1:750 dilution in 2.5% (w/v) BSA in phosphate-buffered saline [PBS; 140 mM NaCl, 2.7 mM KCl, 6.5 mM Na_2_HPO_4_ × 2H_2_O, 1.5 mM KH_2_PO_4_, pH 7.3] for SIMK localization and with mouse anti-phospho-p44/42 primary antibody at 1:400 dilution in 2.5% (w/v) BSA in PBS to visualize activated MAPKs. To improve antibody penetration, vacuum infiltration was used (3×5 min), followed by overnight incubation at 4°C. Samples were then sequentially incubated with appropriate Alexa Fluor-conjugated secondary antibodies. First, Alexa Fluor 647 rabbit anti-mouse secondary antibody (Abcam) diluted 1:500 in 2.5% (w/v) BSA in PBS was used for 2h incubation at 37°C. Samples were extensively washed in PBS (5×10 min), blocked in 5% (w/v) BSA in PBS for 20 min at RT, and incubated with Alexa Fluor 555 goat anti-rabbit secondary antibody (Abcam) by keeping the same dilution and incubation conditions. Nuclei and *S. meliloti* were visualized with 1 μg·ml^-1^ DAPI diluted 1:1000 in PBS for 15 min at RT in darkness.

### FM4-64 staining

The fixable variant of the styryl dye FM4-64 (FX) was used for *in situ* visualizations of plasma membranes in alfalfa root cells treated with *S. meliloti*. Roots were labeled in liquid FAH-N_2_ medium (pH 6.5) containing FM4-64 (FX) at a final concentration of 4 μM in 5 ml Eppendorf tubes for 20 min. The whole labeling was performed on ice. The excess dye was quickly washed out with liquid FAH-N_2_ medium, roots were cut into 1.5 cm long segments and immediately fixed as described previously (Tichá *et al*., 2020). Fixed root segments were used for immunolabeling as described above. For SIMK immunostaining in FM4-64 (FX)-labeled samples, rabbit SIMK-specific primary and Alexa Fluor 647 goat anti-rabbit secondary antibodies were used.

### Confocal laser scanning microscopy (CLSM) and Airyscan CLSM

Root samples immunolabeled for SIMK and activated MAPKs were mounted in the antifade mounting medium [0.1% (w/v) paraphenylenediamine in 90% (v/v) glycerol buffered with 10% (v/v) PBS at pH 8.2 - 8.6] to protect samples from photo-bleaching and used for microscopy. Imaging of immunolabeled SIMK and activated MAPKs was performed with Zeiss LSM 710 platform (Carl Zeiss, Germany) equipped with Plan-Apochromat 40×/1.4 Oil DIC M27 and Plan-Apochromat 63×/1.4 Oil DIC M27 objectives. Samples were imaged with excitation laser lines at 405 nm for DAPI, 488 nm for detection of GFP, 561 nm for Alexa Fluor 555 to visualize SIMK and 631 nm for Alexa Fluor 647 to detect activated MAPKs. Microscopic analysis of immunostained SIMK and FM4-64 FX-visualized membranes in rhizobia-infected root hairs was performed with a Zeiss LSM 880 Airyscan equipped with 32 GaAsP detectors (Carl Zeiss, Germany) using a Plan-Apochromat 63×/1.4 Oil DIC M27 objective. Samples were imaged with excitation laser lines at 561 nm for FM4-64FX and 631 nm for Alexa Fluor 647.

### Quantification of ITs

For quantitative evaluation of ITs formation, 18 days old plants of RSY, *SIMKK-RNAi* and GFP-SIMK lines, originating from somatic embryos, were inoculated with *S. meliloti* wild-type. Inoculated plants were daily subjected to microscopic observations, ITs were counted, and evaluation of ITs number per the whole root system length was performed at 10 dpi with *S. meliloti* using Axio Zoom.V16 Stereo microscope (Carl Zeiss, Germany).

### Image acquisition and processing

The image acquisition, post-processing, semi-quantitative profile measurements, quantitative colocalization analysis, maximum intensity projections from individual z-stacks, subset creation of all fluorescence images, and 3D modeling was made using Zeiss ZEN software (Black and Blue versions, Carl Zeiss, Germany). Data obtained by LSFM imaging were subjected to 3D rendering. A subset of selected z-stacks was created from the whole root volume to capture different stages of root nodulation. Data were imported to Arivis Vision4D 2.12.6 software (Arivis AG, Rostock, Germany), automatically converted to a *.sis file, displayed as 3D objects, and rendered in the maximum intensity mode. Animations and videos were prepared by clipping 3D models in XZ and YZ planes and by using rotation and zooming tools in the 4D clipping panel by arranging keyframes. Although quantification of fluorescence intensities is not influenced by post-acquisition look-up table (LUT) intensity adjustments, all images used for semi-quantitative and quantitative analyses were acquired at the same imaging conditions. The same laser attenuation values for all laser lines were set prior to the acquisition and the thickness of individual optical sections was optimized according to Nyquist criteria. The pinhole sizes for green (GFP), red (Alexa Fluor 555) and yellow (Alexa Fluor 647) channels were matched and the range of detection was appropriately adjusted to ensure separation of emission wavelengths and to prevent fluorescence spectral bleed-through. The brightness and contrast of all acquired images were uniformly adjusted and images exported from ZEN software were assembled into final figure plates using Microsoft PowerPoint.

### Semi-quantitative analysis of the fluorescence intensity distribution

Data obtained from LSFM imaging were semi-quantitatively evaluated by profile measurements to study the fluorescence intensity distribution of GFP-tagged SIMK in alfalfa root hairs and its association with *S. meliloti* at early symbiotic stages. GFP-SIMK mean fluorescence intensity was quantitatively evaluated in specific regions of interest (ROIs) inside root hairs growing under control conditions or interacting with rhizobia. Distribution of SIMK, GFP and activated MAPKs around early invasion structures was determined on fixed and immunolabeled samples inside root hairs of alfalfa control and transgenic plants by semi-quantitative analysis and profile measurement of fluorescence intensities. Intensity profiles were quantified across infection pockets and ITs as indicated in appropriate images. These analyses were done using a profile or measure function of Zeiss ZEN 2011 software (Black version) from single confocal optical sections or maximum intensity projections.

### Quantitative colocalization analysis

The mode of fluorescence signals colocalization was analyzed on immunolabeled root samples of alfalfa control and transgenic plants co-cultivated with *S. meliloti*. Quantitative colocalization analysis between SIMK and activated MAPKs was conducted in particular ROIs at early symbiotic stages around infection pockets and ITs. The colocalization range was measured from single plane confocal sections, and in total, three independent optical sections per infection pocket and IT were analyzed using the colocalization tool of Zeiss ZEN 2014 software (Blue version). Background thresholds were automatically implemented by the iterative Costes approach (Costes *et al*., 2004) and colocalization data were calculated from manually selected ROIs. Data were displayed in intensity-corrected scatterplot diagrams, the intensity correlation of colocalizing pixels was expressed by Pearson’s correlative coefficient and results were graphically edited using Microsoft Excel.

### Statistical analysis

Statistical parameters of performed experiments, number of samples (N), type of statistical test and statistical significance represented by asterisks are included in the figure legends. The statistical significance of differences between treatments was calculated in Microsoft Excel using a t-test and it is indicated by asterisks (* p < 0.05, ** p < 0.01, *** p < 0.001, n.s. no statistical significance).

## Supplemental Data

**Supplemental Figure S1**. Quantitative colocalization analysis of MAPKs around infection pockets in root hairs of control and transgenic plants during an early stage of *M. sativa* – *S. meliloti* symbiotic interaction.

**Supplemental Figure S2**. Quantitative colocalization analysis of MAPKs around ITs in root hairs of control and transgenic plants during *M. sativa* – *S. meliloti* symbiotic interaction.

**Supplemental Figure S3**. Effectivity of ITs formation in control and transgenic plants after inoculation with *S. meliloti*.

**Supplemental Movie S1**. 3D volumetric root rendering of GFP-SIMK line symbiotically interacting with *S. meliloti* expressing mRFP.

**Supplemental Movie S2**. Orthogonal projection of the root hair showing GFP-SIMK association with rhizobia at the docking site from a X-Z view.

**Supplemental Movie S3**. Orthogonal projection of the root hair showing GFP-SIMK association with rhizobia at the docking site from a Y-Z view.

**Supplemental Movie S4**. Orthogonal projection of the root hair showing GFP-SIMK association with a cluster of rhizobia located at the infection site in the neck of root hair curl from a X-Z view.

**Supplemental Movie S5**. Orthogonal projection of the root hair showing GFP-SIMK association with a cluster of rhizobia located at the infection site in the neck of root hair curl from a Y-Z view.

**Supplemental Movie S6**. Orthogonal projection of the root hair showing a very tight association of GFP-SIMK with rhizobia at the infection site before rhizobia entry from a X-Z view.

**Supplemental Movie S7**. Orthogonal projection of the root hair showing a very tight association of GFP-SIMK with rhizobia at the infection site before rhizobia entry from a Y-Z view.

**Supplemental Movie S8**. Orthogonal projection of the root hair showing association of GFP-SIMK with rhizobia entrapped inside root hair curl at the beginning of infection pocket formation from a X-Z view.

**Supplemental Movie S9**. Orthogonal projection of the root hair showing association of GFP-SIMK with rhizobia entrapped inside root hair curl at the beginning of infection pocket formation from a Y-Z view.

**Supplemental Movie S10**. Orthogonal projection of the root hair showing association of GFP-SIMK with rhizobia forming colonies within infection pocket at the beginning of IT formation from a X-Z view.

**Supplemental Movie S11**. Orthogonal projection of the root hair showing association of GFP-SIMK with rhizobia forming colonies within infection pocket at the beginning of IT formation from a Y-Z view.

**Supplemental Movie S12**. Time-lapse imaging of GFP-SIMK accumulation in the nucleus and at the infection site in the root hair during rhizobia attachment.

**Supplemental Movie S13**. Time-lapse imaging of GFP-SIMK accumulation around infection pockets in the root hair.

**Supplemental Movie S14**. Time-lapse imaging of GFP-SIMK accumulation in the nucleus and at the infection site in the root hair during rhizobia attachment analyzed by semi-quantitative GFP-SIMK fluorescence intensity distribution.

**Supplemental Movie S15**. Time-lapse imaging of GFP-SIMK accumulation around infection pockets in the root hair analyzed by semi-quantitative GFP-SIMK fluorescence intensity distribution.

## Acknowledgment

We would like to thank Katarína Takáčová and Monika Vadovičová for their technical help in all stages of the presented work and Dr. Pavlína Mikulková for her expert technical help with LSFM imaging.

## Authors’ contributions

KH, OŠ and MH conducted immunolocalization experiments, KH, MO and OŠ conducted CLSM, ACLSM and LSFM microscopic documentation, KH performed the quantitative evaluation and statistical analyses. MH prepared and selected transgenic alfalfa lines. JŠ and MO contributed to the experimental plan and data interpretation. KH and MO wrote the manuscript with input from all co-authors. JŠ provided infrastructure and secured funding.

## Data Availability

Data that support the findings of this study are available from the corresponding author upon reasonable request.

